# A genome-wide segmentation approach for the detection of selection footprints

**DOI:** 10.1101/2023.11.22.568282

**Authors:** Tristan Mary-Huard, Guillem Rigaill

## Abstract

**Motivation:** In population genetics, the detection of genomic regions under positive selection is essential to understand the genetic basis of locally adaptive trait variation. We propose a principled approach to detect those regions that combines a robust moment based *F*_*ST*_ estimator with a segmentation algorithm.

**Results:** Our approach allows for pairwise comparisons of populations and does not require any prior knowledge about the size of the regions to be detected. The procedure runs within seconds even for large genome datasets with millions of SNPs, and provides a complete landscape of the *F*_*ST*_ distribution over the chromosome. The procedure comes with a grounded estimator of the baseline *F*_*ST*_ level, allowing the detection of regions exhibiting high departures from this reference value. The potential of our procedure is illustrated in two applications in animal and human population genetics. We were able to recover in a matter of seconds regions known to be under selection, often with greater precision than what was reported in previous studies.

**Availability:** Our approach is implemented in the `fst4pg` R package available from the CRAN repository. The Sheep dataset is downloadable from the Zenodo repository https://doi.org/10.5281/zenodo.237116. The 1000 Genome dataset is downloadable from ftp.1000genomes.ebi.ac.uk/vol1/ftp/release/20130502

## 1 Introduction

In population genetics, the detection of loci under positive selection is essential to understand the genetic basis of locally adaptive trait variation. Several methods have been proposed to identify loci subject to local adaptation, see Foll and Gaggiotti (2008); Duforet-Frebourg et al. (2014) and many others reviewed in Hoban et al. (2016). From a methodological point of view, the by-default approach consists in measuring local genetic distances between each pair of populations through the *F*_*ST*_ index (Weir and Cockerham, 1984; Weir and Hill, 2002). One then scans the genome for regions exhibiting an *F*_*ST*_ value significantly higher than the baseline chromosome level. With a few exceptions (e.g. Fariello et al., 2013) the local analysis is performed at the marker level (Foll and Gaggiotti, 2008; De Villemereuil and Gaggiotti, 2015; Duforet-Frebourg et al., 2016).

Although the previously described strategy has led to the identification of many genomic regions under selection in animal (Gouveia et al., 2014; Zhao et al., 2015) plant (Cavanagh et al., 2013) and human (Akey et al., 2002; Duforet-Frebourg et al., 2014) applications, performing the detection at the marker level represents a major limitation in terms of statistical power. Indeed, selection events are expected to affect regions of the genome tagged with several successive SNPs rather than one SNP due to linkage disequilibrium. Note that the region is likely to be larger for recent selection events. A popular solution to bypass this limitation (Hoban et al., 2016; da Silva Ribeiro et al., 2022b) is to perform the analysis through a window-based procedure, using a user-defined window size. As discussed in Hoban et al. (2016) the choice of the window size is critical. From a statistical point of view, the window size should depend on the sizes of the regions to be detected (see Fryzlewicz (2014) Figure 2 for an illustration). A too small window size results in a loss of detection power as one does not consider all of the relevant SNPs. Alternatively a too large window size also results in a power loss since the amplitude of the selection signal is shrinked by agregating selected and non-selected SNPs. Furthermore, the choice of the summary statistic to be used within the window is still under debate (da Silva Ribeiro et al., 2022a).

In this paper, we reformulate the problem of detecting regions with abnormally high *F*_*ST*_ levels as a multiple changepoint detection or segmentation problem (see Fearnhead and Rigaill (2020) for a short introduction). To this end a two step procedure is proposed: in step 1 the chromosome under study is segmented into regions with homogeneous *F*_*ST*_ levels, and in step 2 regions with unusually high genetic differentiation among populations are identified. The proposed approach has several attractive features, both in terms of computational efficiency and biological interpretation. First, relative to window-based procedures, multiple changepoint approaches do not make any assumption on the size of the segments and have the potential to detect both large and small regions. Second, starting from noisy marker-by-marker *F*_*ST*_ estimates our procedure outputs a complete and interpretable landscape of the *F*_*ST*_ distribution over the chromosome. Third, our procedure relies on statistically grounded and computationally efficient approaches for multiple changepoint detection that have been recently developed (Truong et al., 2020; Fearnhead and Rigaill, 2020).

### Outine of the paper

In section 2 we first establish the connection between moment based *F*_*ST*_ estimation and multiple changepoint detection through the use of a specific weighted quadratic loss function (2.1). We then describe i) how the resulting weighted segmentation problem can be efficiently solved using state-of-the-art tools from the statistical literature (2.2), and ii) our global approach for the identification of regions under selection (2.3). The performance of the approach is then illustrated in section 3, both in terms of biological relevancy (3.1) and scalability (3.2). Some elements of discussion are provided in section 4.

## 2 Methods

### 2.1 From *F*_*ST*_ estimators to loss function

In what follows we consider 2 populations from which alleles can be sampled, and we note *p*_*km*_ the reference allele frequency in population *k* at marker *m*. Hereafter we only consider biallelic markers. We first introduce the classical Hudson *F*_*ST*_ along with a moment estimator that will be the starting ground for the choice of the loss function in the segmentation procedure presented in section 2.2.

#### 2.1.1 Hudson *F*_*ST*_

We consider the population analogue of the Hudson estimator introduced in Bhatia et al. (2013), that is based on mismatch probabilities within and between two sampled populations. It is defined as:

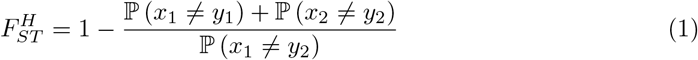

where *x*_*k*_ and *y*_*k*_ are independent indicators for one allele randomly drawn in population *k* = 1, 2. Formally, 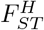 measures the average divergence of Populations 1 and 2 from their most recent common ancestor, see Mary-Huard and Balding (2022) for details.

#### 2.1.2 Moment estimator

Let 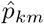 be the empirical reference allele frequency at marker *m*, measured from a sample of *n*_*k*_ gametes drawn at random in population *k*. The Hudson *F*_*ST*_ between populations 1 and 2 can be estimated using the following moment estimator:

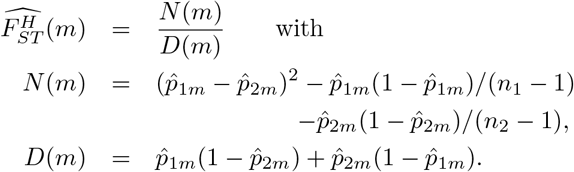

When considering a chromosomic region *R* including several markers the previous estimator can be adapted as follows

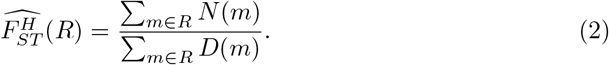

corresponding to the 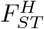 estimator introduced in Bhatia et al. (2013).

#### 2.1.3 Loss function

The moment estimator of equation (2) can be understood as the minimizer of a weighted sum of squares problem. Indeed,

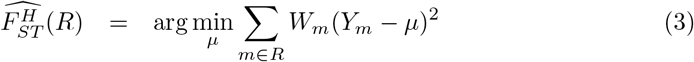

where *Y*_*m*_ = *N* (*m*)*/D*(*m*) and *W*_*m*_ = *D*(*m*).

Recasting the moment estimator as the solution of a (convex) optimization problem has two attractive features. On the one hand expression (3) sheds some light on the robustness of the *F*_*ST*_ moment estimator. From an evolutionary point of view, it can be shown that *D*(*m*) is an unbiased estimator of 2*p*_*m*_(1 − *p*_*m*_), where *p*_*m*_ stands for the reference allele frequency at marker *m* in the most recent common ancestor of Populations 1 and 2 (Mary-Huard and Balding, 2022). The maximum value of *D*(*m*) is reached at *p*_*m*_ = 0.5, corresponding to the most favorable value to evaluate the divergence of the actual frequencies *p*_*m*1_ and *p*_*m*2_ from the ancestral one. Alternatively, if *p*_*m*_ = 0 - i.e. the marker is uninformative - then *D*(*m*) is expected to be close to 0. Therefore using *D*(*m*) as the weight in the optimization problem boils down to weighting the marker contributions according to their (estimated) level of informativeness. On the other hand, interpreting the moment estimator as the minimizer of a least-square loss allows recasting the selective sweep detection task as a multiple change-points detection or segmentation in the mean of the 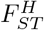 Segmentation is a well-studied problem in the statistical literature and efficient procedures exist and are readily available (Fearnhead and Rigaill, 2020; Truong et al., 2020). Intuitively, we aim to segment the genome in regions of homogenous *F*_*ST*_ (constant weighted mean) and the weighted least square loss provides a natural way to compare any two segmentations. The smaller the loss the better the segmentation.

### 2.2 Weighted least square segmentation of the 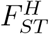 optimization and model selection

From the previous paragraph we argue that chromosomic regions exhibiting differentiated *F*_*ST*_ levels can be identified by segmenting the *F*_*ST*_ signal along the genome using the weighted least square criterion (3). We now present the main steps to define and solve the segmentation problem at hand.

#### 2.2.1 Weighted Least Square segmentation

We consider data *Y*_1_, …*Y*_*n*_ in `R`^*n*^ and weights *W*_1_, …*W*_*n*_. A segmentation *τ* in *D* segments is defined by a set of *D* + 1 ordered changes 0 = *τ*_0_ *< τ*_1_ · · ·, *< τ*_*D*_ = *n* in {0, *n}*. A segment *s* is defined by two consecutive changes : {*τ*_*d*_ + 1,, *τ*_*d*+1_}. For a given number of segments *D* a natural goal is to recover the segmentation that minimizes the following weighted least square (WLS) criterion :

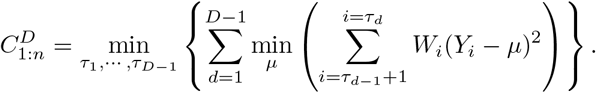

In practice the number and position of the changes are unknown. This raises a statistical problem to select the number of changes and an algorithmic problem to recover for a given number of segments *D* the one realizing 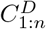 Although procedures have been proposed to handle the unweighted case (Lebarbier, 2005; Rigaill, 2015; Maidstone et al., 2017), they need to be adapted to account for the weights in the loss function and cope with the space complexity as we want to consider large *D*_*max*_ of order 10^2^ − 10^3^ for *n* typically larger than 10^5^. In what follows we present the 3 steps of the procedure that consists in i) computing the WLS criterion 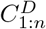 for a set of candidate values *D* ∈ [1, *D*_*max*_], ii) selecting among these candidate values the best one 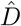 according to a model selection criterion and iii) retrieving the segmentation of size 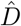 achieving the best WLS criterion.

#### 2.2.2 WLS criterion optimization

For a given *D* the number of possible segmentations in *D* is large: 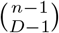 but several efficient dynamic programming algorithms already exist to solve the optimization problem efficiently (Rigaill, 2015; Maidstone et al., 2017; Haynes et al., 2017; Runge et al., 2020). We first adapted the pDPA algorithm of Rigaill (2015) to take into account the weights. As this adaptation was rather straightforward we do not describe it here, but the code is available on the `fpopw` package on the CRAN. The pDPA algorithm recovers 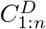 for all *D* in between 1 and a user defined *D*_*max*_ on average in O(*D*_*max*_*n* log(*n*)) time. Its space complexity is in O(*D*_*max*_*n*) which is problematic as we want to consider rather large *D*_*max*_ (i.e. 10^2^ − 10^3^) for *n* typically of order 10^5^ − 10^6^. We describe below a modification of the algorithm reducing space complexity to O(*n*) allowing to handle large *n* and *D*_*max*_.

The pDPA space complexity stems from the storage of two *n* × *D*_*max*_ matrices. The first matrix stores at line *D* and column *i* the best LS solution for a segmentation in *D* of the data from point 1 to *i* : 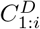 The pDPA algorithm loops over all numbers of segments from 2 to *D*_*max*_. At loop *D* + 1 pDPA only requires rows *D* and *D* + 1 to compute 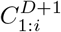 from the 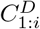 In our implementation, we only store those two lines and the last column that contains the best LS criteria 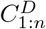 for any number of segments *D* between 1 and *D*_*max*_. This last column will be used in the model selection step to select the number of changes (see next section).

The second matrix stores at line *D* column *i* the best last change for a segmentation in *D* of the data from point 1 to *i*. This second matrix is not used during the dynamic programming step but only at the end to backtrack the optimal segmentation into *D* segments. In our implementation we do not store this matrix, meaning that the optimal set of changepoints 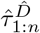 cannot be directly backtracked. This set will be recovered in step iii) as detailed in paragraph 2.2.4.

#### 2.2.3 Model selection

To select the number of segments we use the penalized LS criterion proposed by Lebarbier (2005), that has been shown to have good performances in a recent simulation benchmark (Fearnhead and Rigaill, 2020). The penalty of Lebarbier (2005) has the following form

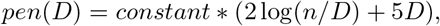

where the constant is related to the variance of the signal and needs to be calibrated from the data. For this calibration we followed the strategy proposed by Lebarbier (2005) and implemented in the `segmentor3isback` R package (Cleynen et al., 2014).

Additionally, we also implemented the penalty proposed by Yao (1988) which is linear in the number of changes: *pen*(*D*) = 2*D* log(*n*)*σ*^2^. As the variance *σ*^2^ is unknown it must be estimated, using e.g. the estimator proposed in Hall et al. (1990). In practice the use of this alternative criterion resulted in an inflated number of segments (results not shown), and we do not considered it further in this paper.

#### 2.2.4 Recovering the selected segmentation

Once 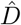 is chosen, we need to retrieve the changepoints : 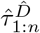 . This can be done using the FPOP algorithm of Maidstone et al. (2017), that minimizes the following LS criterion ^1^:

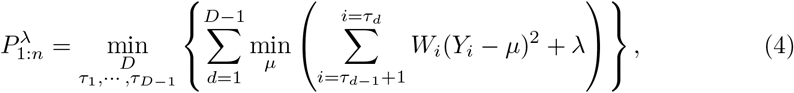

where *λ* is a constant penalty per segment. Following Theorem 3.3 in section 3.2 of Killick et al. (2012) on “Concave penalties” (that covers the case of the penalty of Lebarbier (2005)), if 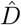 is the best number of segment of a concave penalty *pen*(*D*) then there exists a value of *λ* such that 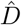 and the corresponding segmentation 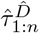 is the solution of (4). In pratice we use FPOP with a penalty *λ* in the interval (*λ*_*min*_, *λ*_*max*_) where

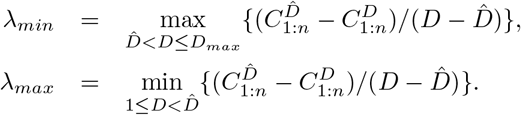

This choice is justified in the following paragraph.

The criteria optimized by FPOP can be rewritten as min 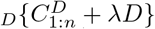. For a *λ*^∗^ in (*λ*_*min*_, *λ*_*max*_) we get, by definition of *λ*_*min*_ and *λ*_*max*_, for any 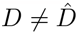 and smaller than *D*_*max*_ that 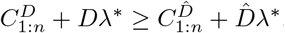. Thus in terms of the FPOP criteria 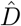 is better than any other *D* in [1, *D*_*max*_]. In theory a value of *D* larger than *D*_*max*_ could be better than 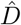 for *λ*^∗^ in (*λ*_*min*_, *λ*_*max*_) but it is our experience that this is not the case for large enough *D*_*max*_.

The time complexity of the FPOP algorithm is on average 𝒪 (*n* log(*n*)). Its space complexity is 𝒪 (*n*). Therefore, not storing the two matrices while running the pDPA and using FPOP to recover the segmentation in 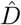 segments yields an average 𝒪 (*D*_*max*_*n* log(*n*)) time and 𝒪 (*n*) space complexity.

### 2.3 Detecting selection footprints through segmentation

#### 2.3.1 Contrasting 2 populations

By definition, the Hudson *F*_*ST*_ defined in (1) contrasts a pair of populations. Consequently, the procedure we propose primarily targets pairwise comparisons between populations. Because the average *F*_*ST*_ level may be quite different from one chromosome to another, the detection of selection sweeps must be performed on each chromosome separately. The output of the segmentation procedure previously detailed applied to the *F*_*ST*_ signal is a serie of adjacent segments and their associated *F*_*ST*_ levels. To detect regions under selection, one needs first to evaluate a reference *F*_*ST*_ level over the chromosome, then identify which segments exhibit a *F*_*ST*_ level much higher than the reference one.

In practice, most if not all *F*_*ST*_ -based procedures require a baseline *F*_*ST*_ level (or some quantities derived from it) as an input. Assuming that selection impacts only a small portion of the markers, we propose to estimate this baseline *F*_*ST*_ level for a given chromosome using

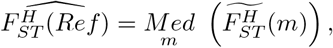

where *Med* corresponds to the median computed over all markers *m* located on the chromosome under study, and 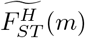 corresponds to the estimated *F*_*ST*_ at marker *m* based on the segmentation, i.e. to the *F*_*ST*_ level of the segment to which *m* belongs to. The resulting estimator is robust as it nullifies the impact of markers belonging to regions associated to outlier *F*_*ST*_ values (i.e. the regions under selection one aims to detect).

Once 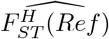 is obtained, one can identify regions under selection by comparing the *F*_*ST*_ level associated with each region with the estimated reference level. In the present paper, we considered a region exhibiting a ratio 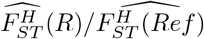 higher than 3 as being under selection.

#### 2.3.2 Contrasting *K* populations

When *K* is larger than 2, the segmentation procedure can be applied pairwise overall possible combinations. Although the number of combinations is 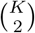, this comes with no prohibitive computational cost since the number of populations is usually low (i.e. *<* 50) and the different segmentations tasks can be parallelized. One then identifies regions under selection by applying the same procedure as described above, then counting for each marker the number of pairwise comparisons for which the ratio is larger than 3. While this approach is arguably very rough from a modeling perspective, we show in the Results section 3.1.3 that in practice the signal of selection may be quite strong so that our rough pairwise approach is sufficient to capture the main regions under selection.

In some applications populations may be gathered into “super-” or “meta-” populations, i.e. subsets of populations defined from prior knowledge. For instance the 1000 Genomes dataset (Consortium et al., 2010) collects information about 26 human populations that have been divided into 5 super-populations (African,Admixed American, East Asian, European and South Asian). One may then be interested in comparing two subsets of populations rather than analyzing the whole set. In such cases the previous procedure can be adapted as follows: rather than considering all pairwise segmentations one can focus on pairwise segmentations contrasting a population from subset 1 with a population from subset 2, then apply the rest of the procedure as previously described. This application case is illustrated in the Results section.

#### 2.3.3 H_1_ enrichment

As mentioned in the Introduction most methods perform selection detection at the marker level. Markers under selection are detected based on a p-value obtained from a test procedure where the *H*_0_ hypothesis corresponds to the absence of detection signal. In contrast our procedure yields a set of detected regions based on the ratio between the local and baseline values of the *F*_*ST*_ . To ease the comparison between the present approach and classical ones we consider an *H*_1_ enrichment criterion. For a givenregion, this criterion corresponds to the proportion of markers with a significant p-value at a given nominal threshold - after correcting p-values for multiple testing. In the following applications multiple testing correction is performed using the Benjamini-Hochberg procedure for FDR control at a nominal level of 10%.

## 3 Results

### 3.1 Sheep dataset

The Sheep dataset includes 507 individuals from 23 domestic sheep populations raised in France, genotyped at 527,823 biallelic markers spread over the 26 autosomal chromosomes. The data are described in Rochus et al. (2018), where a selection sweep detection was performed using the FLK (Bonhomme et al., 2010) and hapFLK (Fariello et al., 2013) methods.

The different aspects of the methodology presented in section 2 are illustrated through 3 different applications: a simple pairwise comparison between 2 populations, a contrast between two sets of populations, and the global analysis of the full dataset.

#### 3.1.1 Two population analysis

We first focus on chromosome 2 that is known to harbor a large region in the vicinity of the myostatin gene (position 109.0-122.3Mbp) that has experienced a large loss of genetic diversity in the Texel population. As a sanity check for our procedure we performed a pairwise comparison between the Texel (TEX) population and the Vendéen (VEN) population that both belong to the Northern Europe group. Figure 1 (left) displays the results of the chromosome 2 segmentation. Each point of the graph corresponds to a marker, located by its genomic position (x-axis) and its *F*_*ST*_ estimate (y-axis). The red broken line represents the segmentation, with the value on the y-axis corresponding to 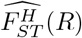 (*R*) for each segment. The resulting profile has 4 segments, two of them (1 and 4) include more than 92% of the markers of the chromosome. Although no constraint is imposed on the *F*_*ST*_ estimates of the different segments, these 2 segments have very similar *F*_*ST*_ levels. Assuming that the regions under selection are small compared to the chromosome size, these 2 segments may be considered as *H*_0_ segments where the baseline *F*_*ST*_ level characterizing the chromosome can be observed. Using the median estimator proposed in section 2.3 yields a baseline *F*_*ST*_ of 0.159, corresponding to the green line on the graph. Assuming a region to be under selection when the *F*_*ST*_ level is at least 3 times higher than the baseline *F*_*ST*_ (red line of the graph) leads to the identification of a region spanning from 108Mb to 122Mb: recovering the known selected region in the vicinity of the myostatin gene.

**Figure 1.**
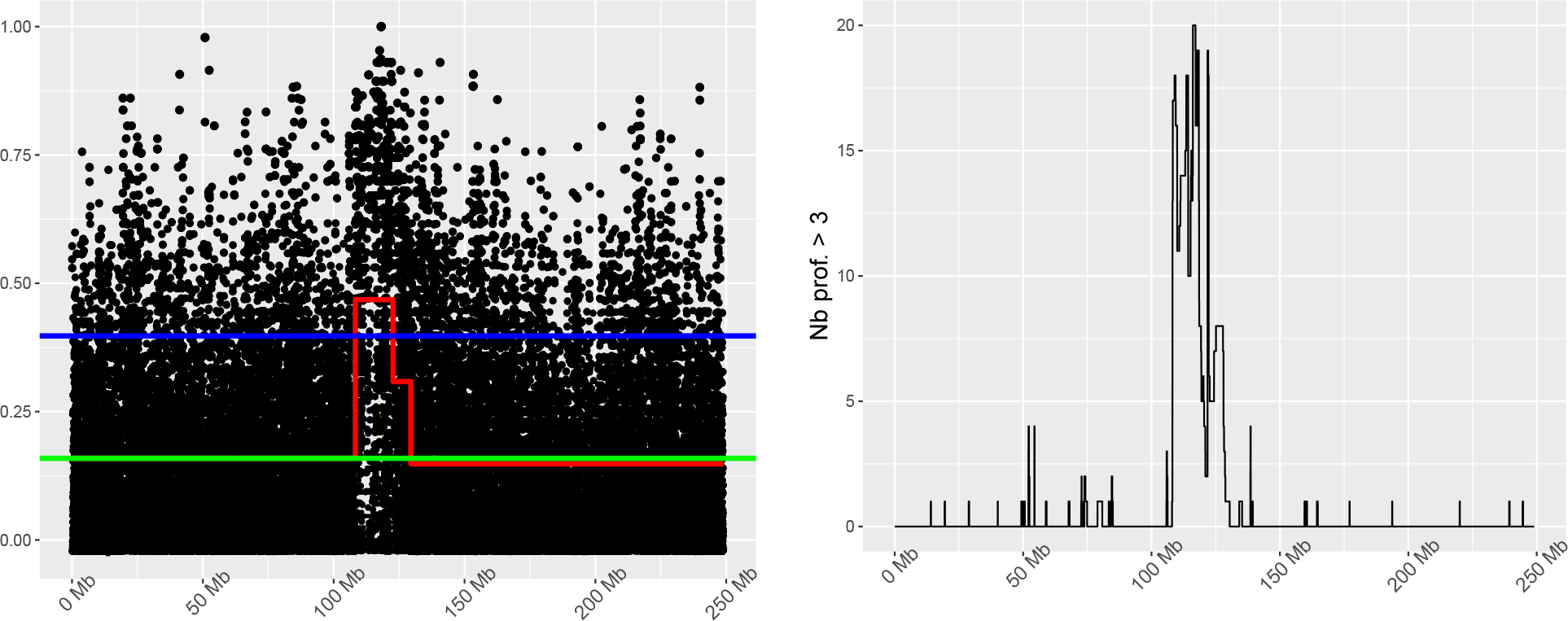
**Left:** Representation of the *F*_*ST*_ distribution between the TEX and VEN populations throughout chromosome 2. Each black point corresponds to a marker, characterized by its genomic position (x-axis) and its associated *F*_*ST*_ estimate (y-axis). The red, green, and blue (broken) lines correspond to the *F*_*ST*_ signal segmentation, the baseline *F*_*ST*_ level, and the chosen *F*_*ST*_ threshold, respectively. **Right:** Number of times a marker was found to have a profiled *F*_*ST*_ value higher than the threshold over all pairwise comparisons, for the same chromosome.

#### 3.1.2 Population subset comparison analysis

To check if the identified region is specific to the TEX-VEN comparison or not, the TEX population is now compared to each of the 21 alternative populations. For each alternative population, a pairwise comparison with TEX is performed as in the previous section, leading to the identification of 21 sets of regions under selection. Figure 1, (right) displays a summary of the 21 analyses: the curve represents the number of times a genomic position has been detected over the 21 comparisons. One can easily identify a major peak spanning from 108Mb to 118Mb detected in at least 10 out of the 21 comparisons corresponding to the known selected region in the vicinity of the myostatin gene.

#### 3.1.3 All pairwise comparison analysis

The global analysis aims at detecting regions under selection without any prior knowledge of the chromosome or population that may be affected. To do so, all pairwise comparisons are performed, then summarized using a graph similar to the one of Figure 1, right. A region was considered under selection if it has been identified in at least 43 of the 253 pairwise comparisons. This threshold corresponds to the theoretical number of pairwise comparisons where an identification would occur if at least 2 out of the 23 populations exhibited a selection pattern in a given region.

A total of 29 regions were found with the present approach. While 23 of them were also found in (Rochus et al., 2018), in each case the region detected with our procedure had narrower boundaries. As an example, the segmentation procedure detected a region on chromosome 14 spanning from 14,1Mb to 14,4Mb that includes the MC1R locus, a gene associated with coat color. This gene was also identified in Rochus et al. (2018), but the length of the reported region was 6 times bigger, spanning from 13,5Mb to 15,2Mb. Similarly, a large genomic region located on chromosome 6 was reported in Fariello et al. (2014) (33.2 to 41Mb using 50K genotypes) and in Rochus et al. (2018) (22.8 to 48.6Mb using the present dataset). This region includes several likely candidate genes: ABCG2 (36,51–36,55Mb) and NCAPG/LCORL (37,25–37,33/ 37,36–37,45Mb) that are respectively associated with milk production, growth and height related traits in several species. Our procedure identified the same set of genes by detecting a region spanning between 34Mb and 39MB, representing a gain of precision of respectively ×1.6 and ×5 compared to the previous analyses.

A complementary *H*_1_ proportion analysis was performed on the basis of the p-values obtained in the initial analysis of Rochus et al. (2018). We used the p-values corresponding to the “north” and “south” FLK analyses (available upon request to the corresponding author of the initial article) as surrogates to p-values for the global analysis. The *H*_1_ proportion was computed for each of the 26 detected regions and ranged from 0 (for regions detected in our analysis but not in the initial article) up to 47%. These high levels confirm that detected regions, while much narrower, focus on chromosomic locations where the selection signal is high.

The 6 regions identified by our procedure and undetected in Rochus et al. (2018) display subsets of populations with highly differentiated frequency patterns (see Suppl. Fig 1). For all regions, the differential pattern strongly overlaps with the Northern/Southern origin of the different populations. Moreover, all detected regions target genic regions, whose corresponding genes have already been associated with traits of interest, see Suppl. Table 1. In particular, the Chr11 region corresponds to the NF1 gene that was identified as a selection signature in a previous study (Fariello et al., 2014). Similarly, The SOX6 gene corresponding to the candidate region on Chr15 is known to be associated with meat quality and body size in sheep, and has already been identified as a putative candidate gene through a genome-wide detection of selective sweep Yang et al. (2016); Zhang et al. (2021).

#### 3.2 1000 Genome Dataset

To illustrate the ability of our procedure to cope with large datasets, we considered the 1000 Genome Dataset Consortium et al. (2010), focusing on the detection of selection sweeps across 2 population subsets: the admixed American populations (CLM, MXL, PEL, PUR) and the European populations (CEU, TSI, FIN, GBR, IBS). We considered 4 chromosomes (1, 2, 15, and 20) representative of the distribution of chromosome size in terms of number of available SNPs. A large number of markers were found to be non-informative for the populations under studies (i.e. with a maximum weight *D*(*m*) over all pairwise comparisons lower than 0.001) and were discarded before the analysis. For a given pairwise comparison regions with an *F*_*ST*_ level higher than 3 times the baseline *F*_*ST*_ were declared under selection, and regions under selection in at least 10 pairwise comparisons (among 20) were reported. This last threshold corresponds to the theoretical number of detections one would expect when at least 2 populations of the smallest subset exhibit a selection pattern compared to the populations of the biggest subset.

Table 1 displays a summary of the results obtained for the different chromosomes. As expected, the estimated number of segments 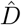 depends on the pair of populations under study, and is not linked to the size (in terms of number of markers) of the chromosome. Figure 2 displays the *F*_*ST*_ profiles on chromosome 15, for the pair of populations with the smallest estimated number of segments 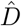 (Pur-TSI) and for the pair with the highest 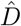 (MXL-GBR). One can observe that whatever 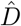 there is a clear baseline *F*_*ST*_ value that can be observed on most of the SNPs through the segmented *F*_*ST*_ profile.

**Table 1.** Number of SNPs before and after filtering, range of the number of segments (over the different pairwise comparisons), number of detected regions and computational time, for each of the 4 chromosomes under study.

| Chr | NbSnp | NbSnp.filt | RangeSegm | NbDetected | Time (s) |
| --- | --- | --- | --- | --- | --- |
| 1 | 6,196,151 | 2,757,470 | 1-42 | 3 | 126 |
| 2 | 6,786,300 | 2,984,399 | 3-40 | 4 | 139 |
| 15 | 2,320,098 | 995,520 | 13-48 | 9 | 32 |
| 20 | 1,739,315 | 772,837 | 5-53 | 6 | 25 |

**Figure 2.**
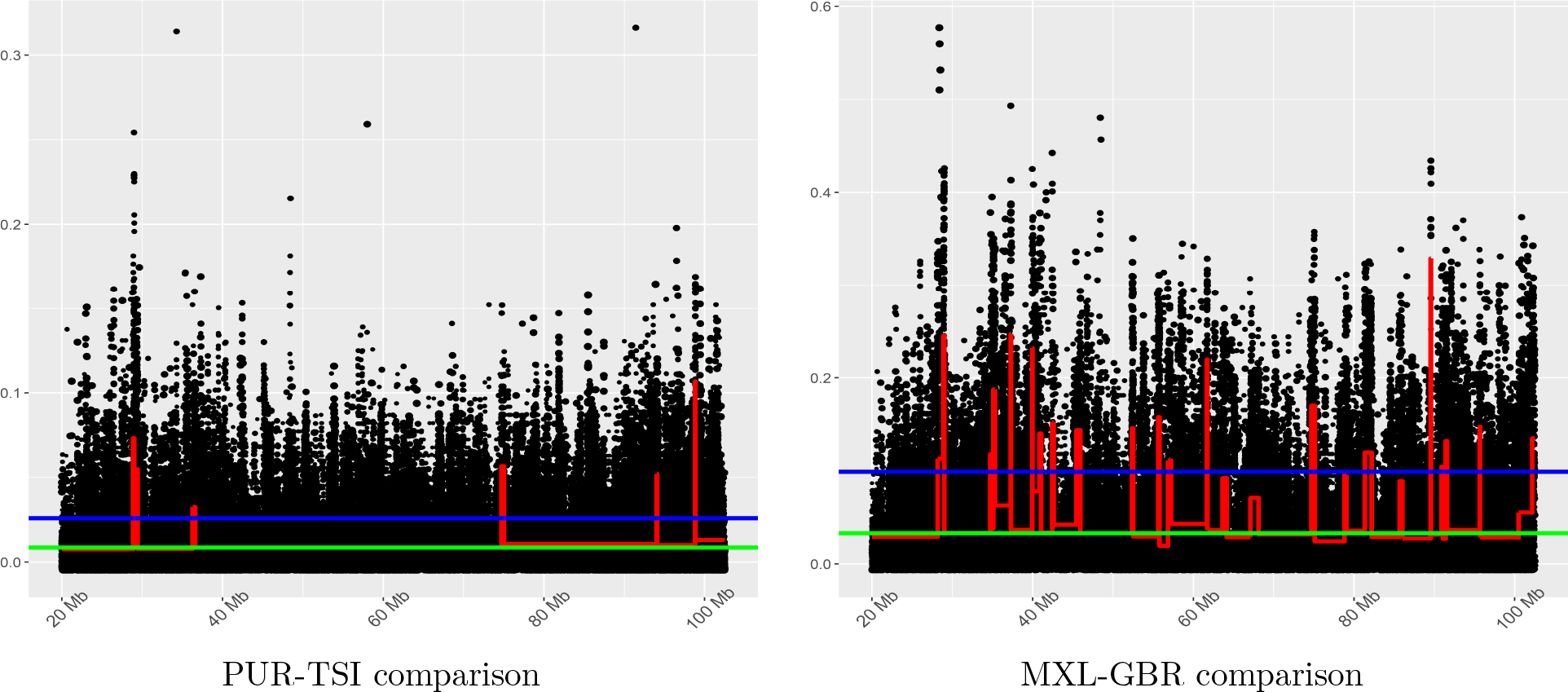
Two *F*_*ST*_ profiles corresponding to chromosome 15, for two different pairwise population comparisons.

Overall 22 regions were detected, with region sizes varying from 4.9kB to 1,1Mb. Beyond a clear differential allele frequency pattern between the European and admixed American populations, many of the identified regions exhibited a strong differential pattern of the PEL population (not shown). As for the previous analysis, most of the identified regions harbored genes that represent likely candidates for selection. For instance, the region spanning from 15,21 to 15,23Mb (including 2,497 SNPs) on chromosome 1 (Suppl. Fig. 2, left) harbours the FLG gene involved in several skin disorders such as atopic dermatitis (Barker et al., 2007), eczema (O’Regan et al., 2008) and ichtyosis vulgaris (Osawa et al., 2011). Similarly, the regions detected at positions 28,17-28,35MB (2,046 SNPs) and positions 79,02-79,04 (327 SNPs) on chromosome 15 (Suppl. Fig. 2, center and right) respectively correspond to the HERC2 and RASGRF1 genes, the first one being involved in human iris color variation (Kayser et al., 2008), and the second one in myopia (Hysi et al., 2010).

## 4 Discussion

In this paper we implemented a segmentation approach coupled with a moment based *F*_*ST*_ estimator to detect genomic regions under positive selection. Our approach allows for pairwise comparisons of populations that can be combined to test for selection across sets of populations. The procedure runs within seconds even for large genome datasets with millions of SNPs, as illustrated on the 1000 Genome application, and is implemented in the `fst4pg` R package available on the CRAN repository.

The applications highlighted the presence of regions under selection that clearly do not reduce to a single SNP, bringing out the need for procedures that do not rely on a single-marker detection. A popular solution to identify regions from a single-marker detection is to combine marker statitics through a region/window-based procedure. However, this strategy has several limitations: the choice of the window size is critical as it should depend on the sizes of the regions to be detected. Another problem is whether to consider overlapping or non-overlapping regions. In the first case, one takes the risk of splitting true regions, and in the second case, one gets highly correlated statistics. The first step of our segementation approach is a statistically grounded Lebarbier (2005); Fearnhead and Rigaill (2020) and computationally efficient Maidstone et al. (2017); Rigaill (2015) solution to these problems as it outputs a reduced set of non-overlapping (small or large) regions whose limits/ends are data-driven. As illustrated in the applications our procedure can detect regions of various sizes without any prior knowledge about the size of the regions to be detected, which represents a major improvement compared to window-based methods.

Regarding the Sheep dataset, our procedure proved to be able to retrieve many of the regions known to be under selection with greater precision in terms of genomic region delimitation compared to previous studies. Although the procedure is based on *F*_*ST*_ and weight estimates that simply rely on the allelic frequency at each marker, it was able to detect regions already known for their haplotypic heterogeneity. This was illustrated by e.g. the detection of the RXFP2 and MC1R genes, which have been previously investigated and are both characterized by multiple adaptive alleles segregating in different breeds of sheep.

A major contribution of our approach is to provide a clear genomic landscape of the *F*_*ST*_ distribution throughout the chromosome. This landscape could not be observed from the pointwise marker estimate representation due to the high variability observed across markers (see Fig 1 and 2). The segmentation also leads to a natural and robust estimation of the *F*_*ST*_ baseline level on each chromosome. This baseline level was used in our applications for the identification of the regions under selection.

In this article, the detection step was performed based on two different criteria: the fold increase for the *F*_*ST*_ compared to the reference level at the population level and the number of pairwise comparisons exhibiting this fold value. The threshold values for these two criteria were chosen to be stringent, as illustrated by the high level of differentiation of the frequency pattern across populations observed for the detected regions. Alternatively, the segmentation procedure could be combined with any test procedure at the region level, such as the ones of Bhatia et al. (2011), Fariello et al. (2013) or Nielsen et al. (2005), or with any test procedure at the marker level using e.g. the *H*_1_ proportion criterion. In such a case the contribution of our procedure will be i) to drastically reduce the computational burden (by providing the test procedure with delimited regions) and ii) to provide robust baseline *F*_*ST*_ estimates to be used as *H*_0_ references, these quantities (or derived ones) being required by most test procedures. While obtaining p-values for any data-driven selected region would be of interest, it remains an open and non-trivial question both for segmentation and sliding windows methods, as emphasized in Hoban et al. (2016) and it is beyond the scope of this paper.

Beyond the detection of selection footprints, providing a full genomic landscape of the *F*_*ST*_ along with delimited regions is likely to be useful for a wider range of applications. In particular we will investigate how the use of regions rather than markers may improve the detection of associations with environmental or phenotypic co-factors (as considered in e.g. environmental GWA analyses, see Fischer et al. (2013)), as demonstrated for GWA Studies in Guinot et al. (2018).

## Acknowledgements

The authors are grateful to Bertrand Servin who contributed to the interpretation of the Sheep dataset.

## Footnotes

1 The adaptation of the FPOP algorithm to the weighted case is straightforward and is not described here.

