## Supplementary Material for "A genome-wide segmentation approach for the detection of selection footprints"

#### 1 Supplementary Figure 1

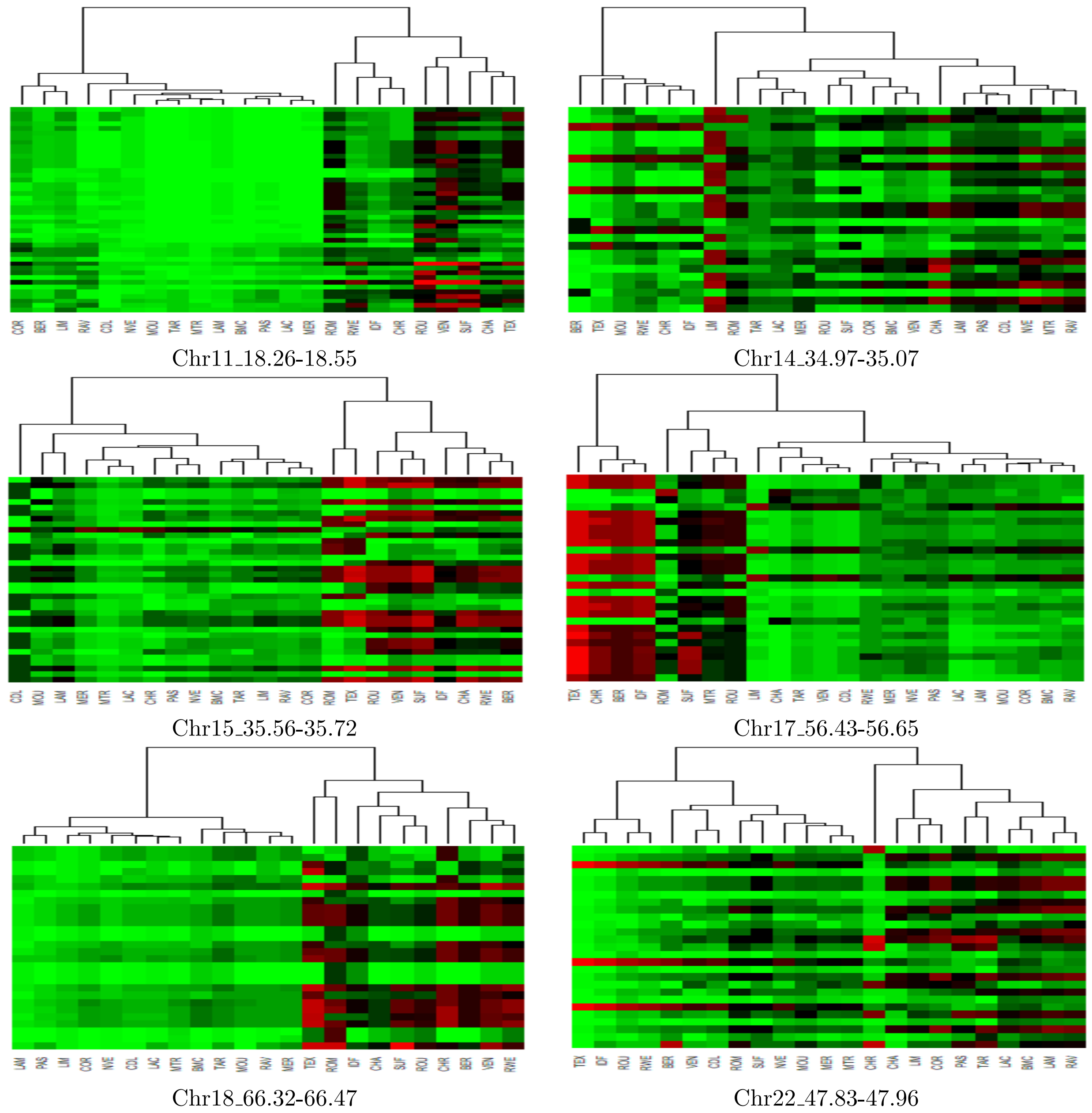


Figure 1: Six regions newly detected with our procedure applied to the Sheep dataset. The $i, j$ entry of the heatmaps corresponds to the alternative allele frequency at marker $i$ (in row) in population $j$ (in column), from green (0) to red (1) through black (0.5).

#### 2 Supplementary Figure 2

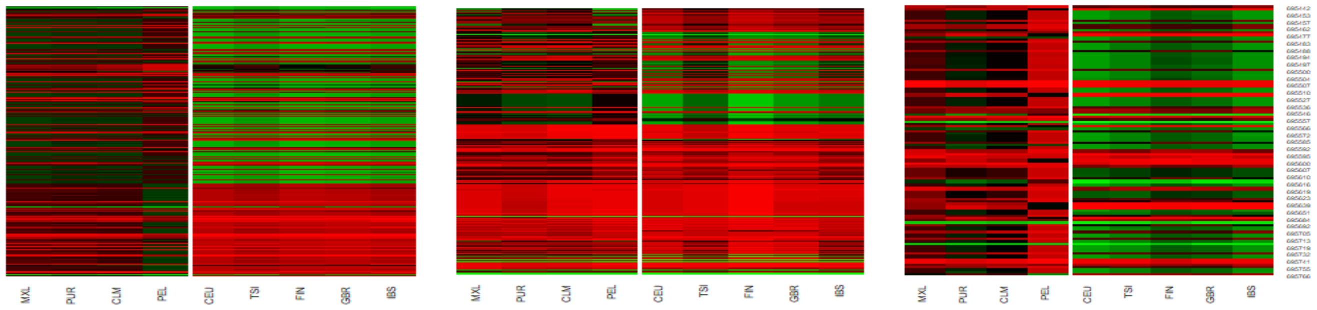


Chr1:15,21:15,23 Chr15:28,17:28,35 Chr15:79,02:79,04

Figure 2: Three regions on chromosomes 1 and 15 identified as candidates to selection in the 1000 Genome dataset. The heatmap representation is similar to the one of Figure 1.

### 3 Supplementary Table 1

| Chr | Start | End | Genes | Ref. |
|---|---|---|---|---|
| 11 | 18268635 | 18557122 | NF1 | [3] |
| 14 | 34974494 | 35076657 | SMPD3, SLCA6OS | [6] |
| | | | | [4] |
| 15 | 35561888 | 35729767 | SOX6 | [8] |
| | | | | [9] |
| 17 | 56438591 | 56656539 | SUDS3, TAOK3 | [7] |
| | | | | [2] |
| 18 | 66326213 | 66470371 | CDC42BPB | [1] |
| 22 | 47835463 | 47961104 | MGMT | [5] |

Table 1: Candidate gene list associated to the 6 regions newly detected with our procedure.
